# Nature and effective range of non-cell autonomous activator and inhibitor peptides specifying plant stomatal patterning

**DOI:** 10.1101/2020.04.29.069211

**Authors:** Scott Zeng, Emily K. W. Lo, Bryna J. Hazelton, Miguel F. Morales, Keiko U. Torii

**Author notes:** Center for Epigenetics, John Hopkins University School of Medicine, Baltimore, MD 21205.

## Abstract

Stomata are epidermal valves that facilitate gas exchange between plants and their environment. Stomatal patterning is regulated by EPIDERMAL PATTERING FACTOR (EPF)-family of secreted peptides: EPF1 enforcing stomatal spacing, whereas EPF-LIKE9, also known as Stomagen, promoting stomatal development. It remains unknown, however, how far these signaling peptides act. Utilizing Cre-Lox recombination-based mosaic sectors that overexpress either EPF1 or Stomagen in Arabidopsis cotyledons, we reveal a range within the epidermis and across the cell layers in which these peptides influence patterns. To quantitatively determine their effective ranges, we developed a computational pipeline, SPACE (Stomata Patterning AutoCorrelation on Epidermis), that describes probabilistic two-dimensional stomatal distributions based upon spatial autocorrelation statistics used in Astrophysics. The SPACE analysis shows that, whereas both peptides act locally, the inhibitor, EPF1, exerts longer-range effects than the activator, Stomagen. Furthermore, local perturbation of stomatal development has little influence on global two-dimensional stomatal patterning. Our findings conclusively demonstrate the nature and extent of EPF peptides as non-cell autonomous local signals and provides a means to quantitatively characterize complex spatial patterns in development.

**Summary Statement:** Non-cell autonomous effects of activator and inhibitor peptides on 2-D spatial patterning of stomata were quantitatively characterized using chimeric sectors and a SPACE computational pipeline.

## Introduction

During the development of multicellular organisms, distinct cell types emerge with specific roles and functions. Cell-to-cell communication of positional cues and spatial information is essential to coordinating the transition from a tissue of uniformly undifferentiated cells into a robust pattern of specialized identities. For plant systems, the presence of the cell wall prevents direct cell-to-cell contact or cell mobility, thereby excluding many of the mechanisms for pattern formation studied in animals, such as the transmembrane receptor Notch and its membrane-bound ligand Delta (Artavanis-Tsakonas et al., 1999), or the contact-dependent depolarization and repulsion between different pigment cell types in zebrafish stripe patterning (Eom and Parichy, 2017; Inaba et al., 2012; Nusslein-Volhard, 2012). The absence of these mechanisms means plants are model systems for isolating and studying the role of local ligand secretion in pattern formation independently of variables such as cell movement or apoptosis (Torii, 2012b).

Stomata, the pores on the plant epidermis responsible for mediating gas exchange and water control, differentiate according to a special cue, which enforces the “one-cell spacing rule” in which no two stomata develop adjacent to each other (Bergmann and Sack, 2007; Pillitteri and Torii, 2012). Locally, stomatal spatial patterning is enforced through a family of small secreted peptide ligands called EPIDERMAL PATTERNING FACTORS (EPFs), which are perceived by a family of ERECTA-family receptor kinases and their signal modulator TOO MANY MOUTHS (TMM) (Hara et al., 2007; Hara et al., 2009; Hunt and Gray, 2009; Lee et al., 2012; Nadeau and Sack, 2002; Shpak et al., 2005; Torii, 2012a). Perception of EPF2 peptide inhibits the entry into stomatal cell lineages (Hara et al., 2009; Hunt and Gray, 2009). Antagonistically, EPF-LIKE9 (EPFL9), also known as Stomagen, is secreted from the subepidermal tissue into the epidermis, and promotes stomatal differentiation via competing for receptor binding with EPF2, and also likely with EPF1 (Kondo et al., 2010; Lee et al., 2015; Sugano et al., 2010). At a later stage, spatial patterning of stomata and differentiation of a stomatal precursor, known as a meristemoid, is controlled by EPF1 (Hara et al., 2007; Qi et al., 2017). Consistent with their function as signaling ligands controlling stomatal development, ectopic overexpression or peptide application of EPF1 and EPF2 confer epidermis devoid of stomata, the former with arrested meristemoids and the latter with reduced stomatal lineage cells (Hara et al., 2007; Hara et al., 2009; Hunt and Gray, 2009). Conversely, Stomagen overexpression or peptide application confers stomatal clusters, resembling the loss of *TMM* or three *ERECTA*-family genes (Kondo et al., 2010; Sugano et al., 2010).

Globally, long range signals are also necessary to optimize stomatal patterning for its physiological functions of mediating gas exchange, water exchange, and photosynthetic efficiency (Hetherington and Woodward, 2003). Small chemical hormones such as ethylene increase stomata (Serna and Fenoll, 1996), whereas others such as abscisic acid reduce their number (Tanaka et al., 2013), but the effect of these individual chemicals can depend on the tissue or species (Qi and Torii, 2018). Auxin is another hormone that broadly regulates plant development, but its inhibition of stomatal density partly depends on the absence of light, illustrating the integration of environmental information as another set of signals (Balcerowicz et al., 2014; Hronkova et al., 2015; Zhang et al., 2014). Furthermore, environmental factors perceived in mature leaves may affect density in younger leaves, demonstrating a spatial propagation of signaling that connects local and global contexts of patterning (Casson and Gray, 2008).

Whereas endogenous and environmental factors controlling stomatal development have been described in detail, much less well understood is how these signals propagate their efficacy in cell-to-cell communication to constitute the emergence of stomatal spatial patterning across the epidermis. The expression of Stomagen in the mesophyll indicates non-cell-autonomous effects across tissue layers, but the range of these signals as they travel between these cells is unknown. One way to assess the movement of signaling peptides is to directly visualize their movement. However, the addition of fluorescent protein tags, such as GFP, impairs the movement of peptide hormones, and the highly processed nature of some peptides hampers such approach. Moreover, such visualization does not address the extent of how signaling peptides influence the local spatial patterning of stomata or whether there is any intersection with global epidermal patterning.

To address these questions, we harnessed Cre-*/ox* recombination and the GAL4/UAS transactivation system (Heidstra et al., 2004) to generate mosaics in which peptide overexpression was localized to sectors of epidermal tissue. To quantitatively analyze these effective ranges, we then developed SPACE (Stomata Patterning Auto Correlation on Epidermis), a computational pipeline that applies spatial correlation techniques. Rather than traditional stomatal phenotype metrics, such as stomata index or density, neither of which describes the two-dimensional spatial patterning, our SPACE analysis revealed the effective range of EPF and Stomagen peptides in influencing epidermal patterning. Our study establishes the roles of EPF-family peptides as signals for cell-to-cell communication and the ranges at which they act. Our study also highlights the use of a spatial correlation approach to analyzing stomata patterning that can be adapted for analyzing both local and global signals, addressing the growing need for such techniques in phenotypic analysis of pattern formation.

## Results

### Genetic Mosaic Analysis Demonstrate Non-Cell-Autonomous actions of EPF1 and STOMAGEN

To address how EPF/EPFL peptides spatially influence stomatal patterning in a non-cell autonomous manner, we generated seedlings with genetic mosaic sectors overproducing individual EPF/EPFL peptides of opposite biological functions: EPF1, which restricts stomatal development and STOMAGEN, which promotes stomatal development (Hara et al., 2007; Sugano et al., 2010). For this purpose, we implemented Cre-*/ox* recombination coupled with the two-component GAL4/UAS transactivation system (Heidstra et al., 2004). Here, heat-shock treatment induces the expression of a CRE recombinase, which acts on two Lox-p sites to create GAL4+ sectors. Within the sectors, both endoplasmic reticulum-trapped green fluorescent protein (GFP_ER_), which marks the sectors in a cell autonomous manner, and *EPF/EPFL* peptide genes (either *EPF1* or *STOMAGEN/EPFL9*) were simultaneously overexpressed (Fig. 1A, B). To accurately monitor the non-cell autonomous effects of these *EPF/EPFL* genes, we expressed the non-epitope tagged EPF1 and STOMAGEN rather than a fluorescent proteins fusion (e.g. CFP/RFP) that may impact the behavior of these small secreted peptides. Our heat-shock conditions yielded high frequency of genetic mosaics per seedlings screened (13.9% to 100%; Table S1). Durations of heat-shock treatment were carefully analyzed to yield sectors of comparable size and number per cotyledon (see Methods). Quantitative RT-PCR analysis confirmed that our heat-sock treatment led to elevated expressions of *EPF1* and *STOMAGEN* transcripts (Fig. 1C).

**Figure 1.**
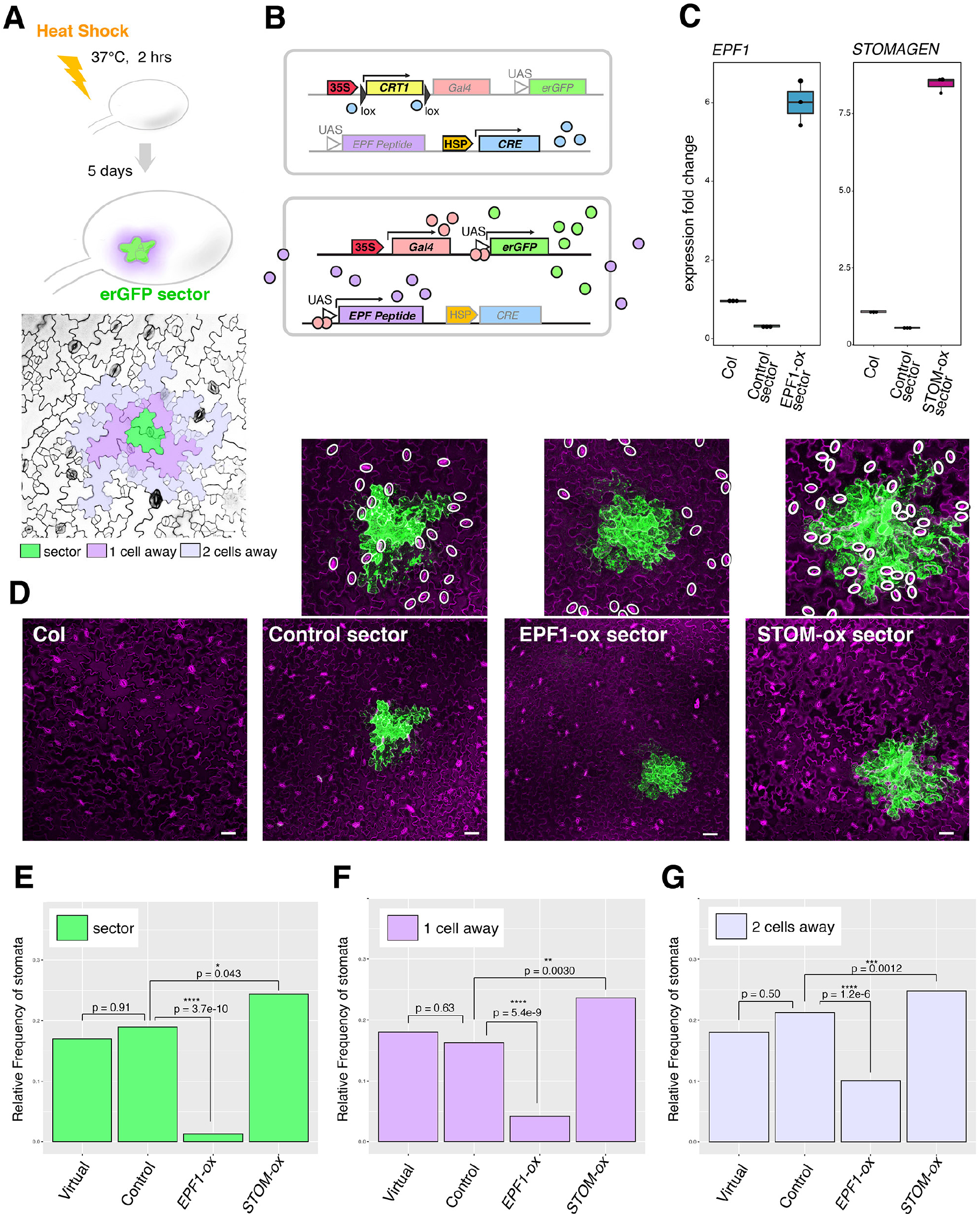
Mosaic Sectors Overexpressing EPF Peptides Non-cell Autonomously Influence Stomatal Patterning. (A) Schematic diagram of the experimental design to generate erGFP sectors by heat-shock treatment (top); False-colored confocal microscopy image of an abaxial epidermis. Green, a sector; Purple, cells immediately adjacent to the sector (1 cell away); Lilac, cells 2 cells away from the sector (B) Schematic diagram of heat-shock induced Cre-Lox recombination and induction of endoplasmic-reticulum trapped GFP (erGFP) as well as secreted EPF peptides. (C) Quantitative RT-PCR analysis of transcripts of *EPF1* (left) and *STOMAGEN* (right) from 7-day-old seedlings of non-transformed Col, control sector expressing erGFP only and sectors expressing *EPF1* or *STOMAGEN*. The transcripts are normalized against Actin (*ACT2*). Three biological replicates were performed, each with three technical replicates, and representative results are shown. (D) Z-stacked, tile-scanned representative confocal microscopy images of 7-day-old cotyledons subjected to heat-shock treatment as described in (A). From left, non-transformed Col, transgenic lines expressing a control sector, EPF1 overexpressing sector, and STOMSGEN overexpressing sector. Scale bars, 50 μm. Above insets, Close-up images of each sector with stomata highlighted by white ovals. For tile scan of the entire cotyledons, see Fig. S1. (E-G) Relative frequency of stomata (number of stomata per total number of epidermal cells) within sectors (E; green), cells immediately adjacent to sectors (F; purple, “1 cell away”), and cells adjacent to immediate neighboring cells (G; lilac, “2 cells away”). For wild type, virtual sectors of the same size and geometry as real sectors were computationally placed. Total numbers of stomata and epidermal cells were aggregated to generate a single dataset for each genotype to enable robust statistical testing. Number of sectors subjected to analysis; Number of sectors subjected to analysis; n=20 (virtual), n=34 (Control), n=25 (EPF1), n=31 (Stomagen). Total number of stomata and non-stomatal epidermal cells counted in sectors; n=229 (virtual), n=364 (Control), n=228 (EPF1), n=410 (Stomagen). Total number of stomata and non-stomatal epidermal cells counted adjacent to sectors; n=384 (virtual), n=564 (Control), n=358 (EPF1), n=817 (Stomagen). Total number of stomata and non-stomatal epidermal cells counted adjacent to immediate neighboring cells; n=702 (virtual), n=952 (Control), n=633 (EPF1), n=1398 (Stomagen). A χ-square analysis was performed to test significant deviation between the frequencies of two aggregated samples.**, p<0.05; ***, p<0.005; ****, p<0.0005.

To test that GFP_ER_ expression alone would not affect stomatal patterning or density, we also heat-shock treated seedlings harboring control empty-vector to generate control GFP sectors that do not overexpress EPF1/STOMAGEN peptide (Fig. 1). Furthermore, to determine whether the shape of sectors itself would affect quantification, sector outlines from mosaics were overlaid onto heat shocked wild-type cotyledons to create virtual “geometric sectors” as another control.

We first analyzed the stomatal phenotype within GFP_ER_-marked sectors (Fig. 1C, D). As expected, stomatal index (SI: number of stomata/(number of stomata + non-stomatal epidermal cells) x100) was significantly reduced within the EPF1-expressing sectors (p=3.7e-10) whereas it increased within the STOMAGEN-expressing sectors (p=0.043) (Fig. 1D). No statistical difference was observed in the stomatal index within control empty sectors when compared to the geometric sectors on wild type (p=0.91) (Fig. 1D), confirming that heat shock treatment or GFP_ER_ expression does not influence stomatal development, and that sector shape does not bias quantification. Within the STOMAGEN-sectors, stomata developed in clusters (Fig. 1C), thus verifying that our sector overexpression functioned as intended (Hara et al., 2007; Kondo et al., 2010; Lee et al., 2015; Lee et al., 2012; Sugano et al., 2010).

Next, we examined whether stomatal development in epidermal tissue near but not inside the GFP_ER_ sectors was also inhibited or promoted by peptide overexpression from these sectors (Fig. 1A, B, D-G). Examining the confocal images of cotyledon epidermis, it appears that regions surrounding EPF1-sectors tend to be devoid of stomata, whereas regions surrounding STOMAGEN-sectors differentiate more stomata (Fig. 1D). For a quantitative analysis, we measured the stomatal index for cells within each sector (Fig. 1A, bottom panel, green), adjacent to a sector (Fig. 1A, bottom panel, purple), or neighboring a sector-adjacent cell (Fig. 1A, bottom panel lilac). Indeed, stomatal index of cells adjacent to EPF1-expressing sectors was reduced (p=5.4e-9), and stomatal index near STOMAGEN-expressing sectors was increased (p=0.0030) (Fig. 1F). On the other hand, stomatal index adjacent to control empty sectors was not statistically different from those inside the control control empty sectors or on heat-shocked wild-type geometric sectors (Fig. 1E, F). Likewise, the stomatal index on cells that neighbored a sector-adjacent cell, i.e. those at a two-cell distance away from the sector (Fig. 1G). It can be seen, however, that whereas the stomatal index near EPF1-expressing sectors remained lower than in control empty vectors or wild-type, stomatal production gradually increased for cells farther away from EPF1-sectors (Fig. 1G), suggesting that sector’s impact on stomatal patterning weakened with distance. Combined, these mosaic sector analyses directly demonstrate the non-cell autonomous actions of EPF/EPFL peptides in adjacent and nearby epidermal cells.

### EPF1 Secreted from the Mesophyll Can Inhibit Stomatal Development

Stomagen is known to secrete from the developing mesophyll layer to promote stomatal development in the epidermis [4]. To address whether EPF/EPFL family peptides have an intrinsic property to function across tissue layers, we sought to test if EPF1 expressed in the mesophyll could also affect stomatal development in the epidermis. For this purpose, we identified GFP_ER_ sectors induced exclusively in the mesophyll (Fig. 2A-B), and subsequently measured the stomatal index in the adaxial epidermal cells located directly above these sectors (Fig. 2C-E). As expected, Stomagen expression from mesophyll sectors promoted stomatal development (Fig. 2G). Conversely, EPF1 expressing mesophyll sectors inhibited stomatal development, in the adjacent epidermal cells (Figure 2G). As before, we extended our quantification to address whether this disruption to stomatal patterning acted at a larger range, in cells that did not directly neighbor the mesophyll cells of interest (Fig. 2F, H). Taken together, we conclude that, like Stomagen, EPF1 is capable of influencing the epidermis via secretion from the mesophyll in a non-cell-autonomous way if ectopically expressed.

**Figure 2.**
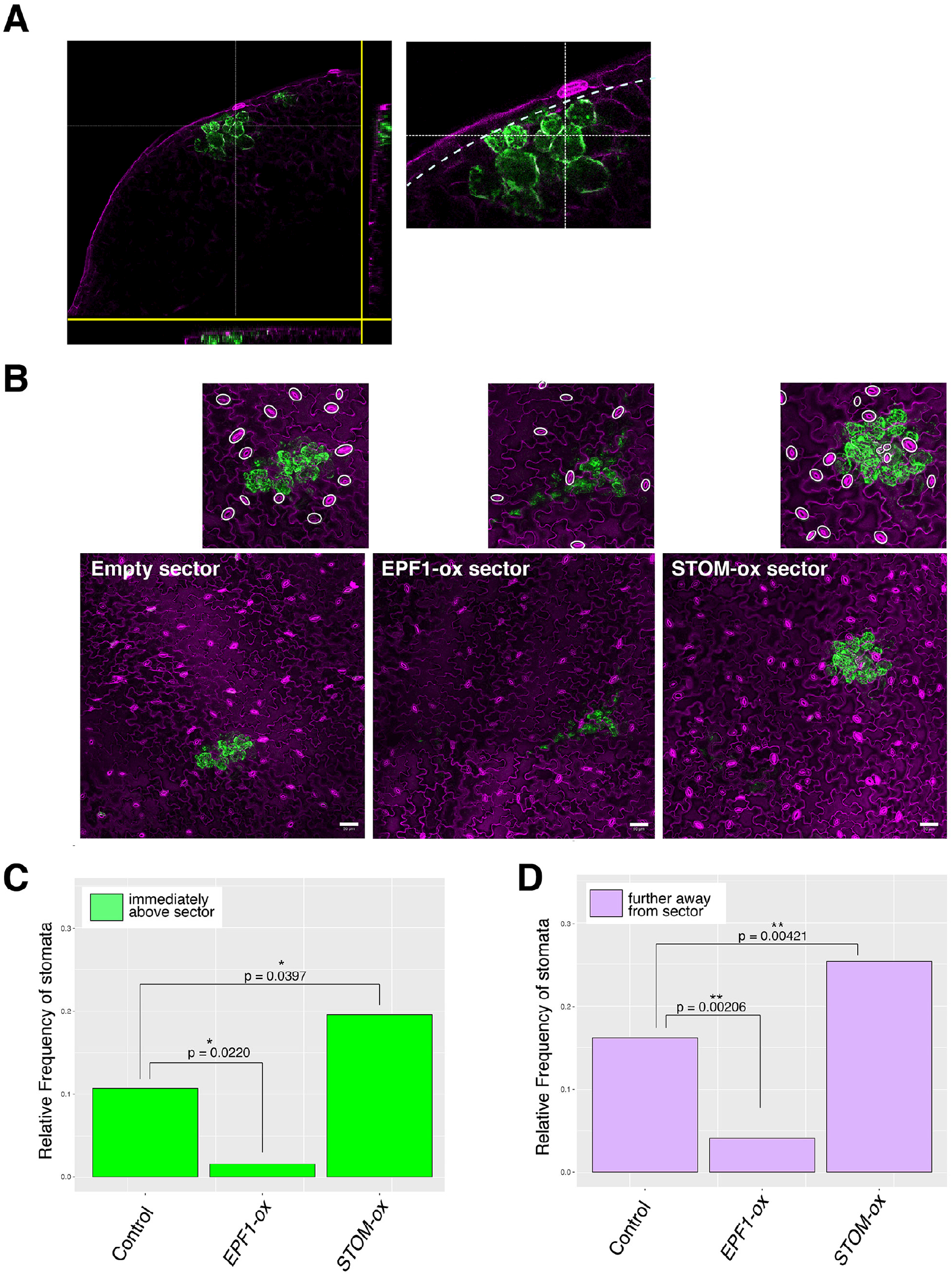
Mesophyll Sectors Overexpressing EPF-family Peptides Locally Influence Stomatal Patterning. (A) Example of Cre-Lox generated mesophyll sector shown as orthogonal slices. Right, close-up of a sector. Dotted line, an inner boundary of epidermal layer. (B) Z-stacked, tile-scanned representative confocal microscopy images of 7-day-old cotyledons with mesophyll sectors. From left, transgenic lines expressing a control sector, EPF1 overexpressing sector, and STOMSGEN overexpressing sector. Scale bars, 50 μm. Above insets, Close-up images of each sector with stomata highlighted by white ovals. For tile scan of the entire cotyledons, see Fig. S2. (C-D) Relative frequency of stomata (number of stomata per total number of epidermal cells) immediately above the mesophyll sector (C; green) and cells immediately adjacent to the cells above the mesophyll sectors (D; purple). Total numbers of stomata and epidermal cells were aggregated to generate a single dataset for each genotype to enable robust statistical testing. Number of sectors subjected to analysis; n=19 (Control), n=8 (EPF1), n=19 (Stomagen). Total number of stomatal and non-stomatal epidermal cells subjected to analysis; n=295 (immediately above the mesophyll sector); n=773 (adjacent to cells immediately above the mesophyll sector). A χ-square analysis was performed to test significant deviation from the stomatal frequency of control plants. **, p<0.05; ***, p<0.005; ****, p<0.0005.

### EPF1 and Stomagen Act in a Limited Effective Range

Our results demonstrate that EPF1 and Stomagen act non-cell-autonomously, but do not address the distance at which these peptides can act to influence epidermal cell fate. We first analyzed this effective range by developing a computational pipeline to quantitatively analyze the stomatal density at various distances relative to the sectors (see Methods). Briefly, full tile-scanned confocal images of entire cotyledons were first converted into a 2-D spatial coordinate plot of the XV-coordinates of every single stomata of an entire cotyledon, sector outlines, and cotyledon outlines (Fig. 3A-C). Subsequently, stomatal density was calculated in the following regions: epidermal tissue inside the GFP_ER_ sector outline (“the sector region”), epidermal tissue located within a 100 μm range of the GFP_ER_ sector outline excluding the sector interior itself (“the nearby region”), and the rest epidermal tissue beyond the 100 μm range (“the faraway region”) (Fig. 3C).

**Figure 3.**
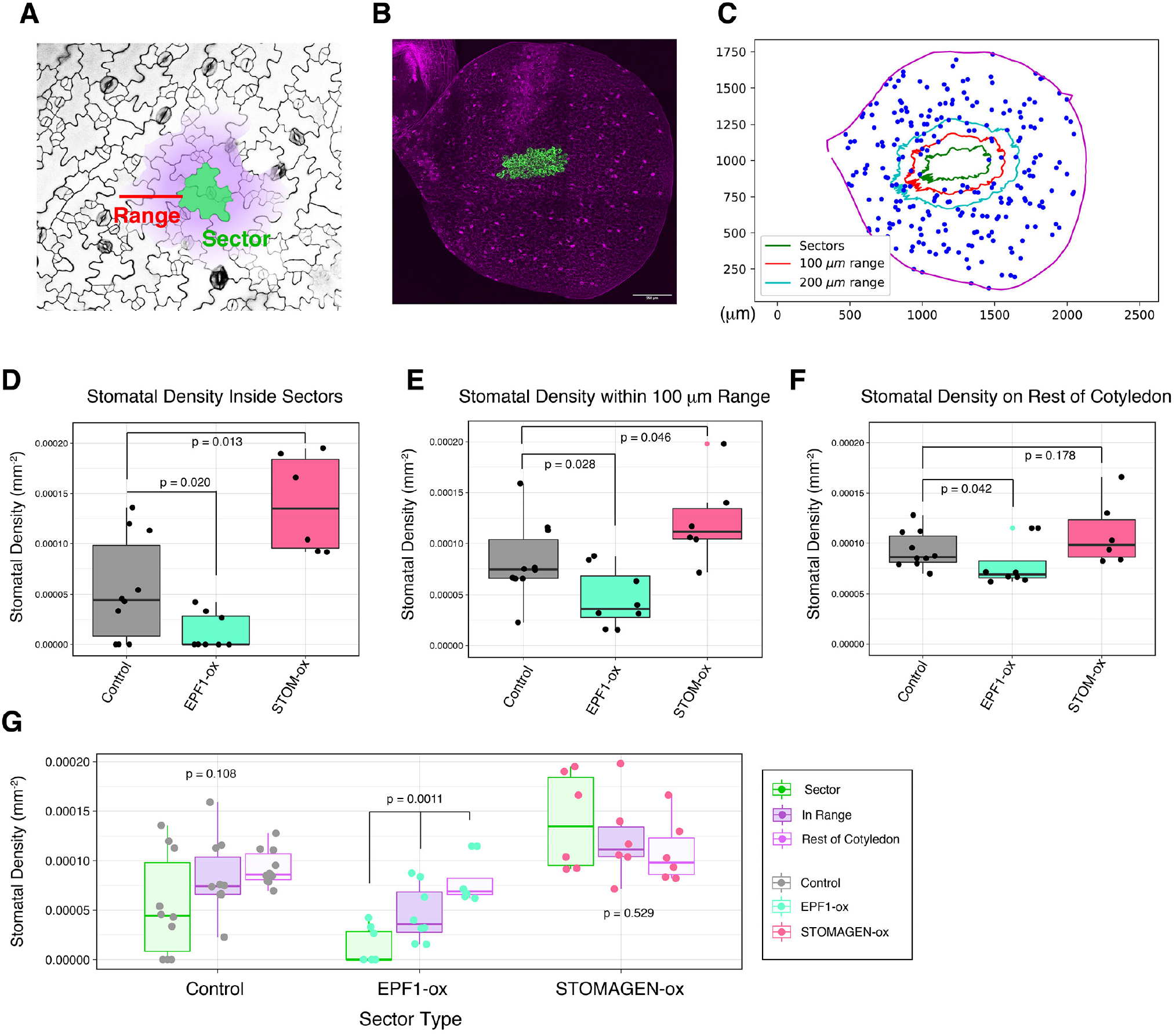
Quantitative Analysis of Effective Range by Non-cell Autonomous Effects of EPF1 and Stomagen-overexpressing Sectors. (A) Schematic diagram. The coordinate outline of a given sector (green) was enlarged to generate a new coordinate boundary at a defined Range (red; in this case, 100 μm) away from the sector outline. The defined range maintains the geometry of the original sector. The stomatal density was calculated in three regions: the interior of the original sector outline (green), the interior of the expanded Range excluding the original sector (purple, but not green); and the rest of the cotyledon (white). (B) Z-stacked, representative tile scan of a whole cotyledon with EPF1-ox sectors. Scale bar, 250 μm. For tile scans of other genotypes, see Figure S1. (C) Representative 2-D coordinate mapping of the cotyledon boundary (magenta), the sector boundaries (green), the boundaries of expanded Ranges at 100 μm (red) and 200 μm, and stomata (blue). Note that if the expanded Range extends beyond the actual cotyledon boundaries, this extended area is excluded from analysis. (D) Stomatal density inside of sectors for individual cotyledons: cotyledons with control sector(s) expressing erGFP only (gray; n=10); cotyledons with sector(s) overexpressing EPF1 (EPF1-ox)(teal; n=8); cotyledons with sector(s) overexpressing Stomagen (STOM-ox)(coral red; n=6). Total number of stomata counted in sectors, n=24 (control); n=5 (EPF1-ox); n=50 (STOM-ox). A Mann-Whitney U test was performed to test significant deviation between distributions of stomatal density. (E) Stomatal density within 100 μm Range of sectors for individual cotyledons: cotyledons analyzed are same as in (D). Total number of stomata counted within 100 μm Range, n=116 (control); n=52 (EPF1-ox); n=148 (STOM-ox). A Mann-Whitney U test was performed to test significant deviation between distributions of stomatal density. (F) Stomatal density outside of the specified regions in (D) and (E) for individual cotyledons: cotyledons analyzed are same as in (D). Total number of stomata counted on the remaining area of cotyledons, n=2229 (control); n=1765 (EPF1-ox); n=1306 (STOM-ox). A Mann-Whitney U test was performed to test significant deviation between distributions of stomatal density. (G) Data provided in (D-F), grouped by sector type and region of stomatal density. For each sector type, a Kruskal-Wallis (non-parametric ANOVA) test was performed to test significant deviation in stomatal density in sectors vs. 100 μm Range vs. rest of cotyledon.

As observed previously via stomatal index, the stomatal density inside EPF1-sectors was reduced and the stomatal density inside STOMAGEN-sectors increased when compared to control, empty vector sectors (p=0.020 for EPF1-sectors; 0.013 for STOMAGEN-sectors)(Fig. 3D). In “the nearby region” of a 100 μm range around sectors, the effect of these peptides on stomatal density remained statistically significant (p=0.028 for EPF1-sectors; p=0.046 for STOMAGEN-sectors)(Fig. 3D, Fig. S2), suggesting that non-cell autonomous actions of these secreted EPF/EPFL peptides are not constrained by cellular geometry. However, STOMAGEN-sectors did not impact the stomatal density of the “faraway region” beyond 100 *μ*m in a statistically significant manner (p=0.178) whereas the presence of EPF1-sectors did (p=0.042), relative to the stomatal density faraway from control sectors (Fig. 3F). The results suggest that the non-cell-autonomous effects of STOMAGEN are local and limited in range. A comparison of within sector, “the nearby region” and the rest of cotyledons within each sector type revealed that both control vector-only sectors and STOMAGEN-sectors do not exhibit statistical significance, whereas a gradual decay of EPF1 effects was evident (Fig. 3G).

We further expanded the “nearby region” to within 200 μm to explore whether doubling the range of would reveal different patterns of effective range (Fig. S3). At this range, the stomatal density of “nearby region” of STOMAGEN-sectors and control empty vectors were no longer statistically differed (p=0.058) whereas that of EPF1 remained effective (p=0.034)(Fig. S3). Similar to the analysis at 100 μm range (Fig. 3), stomatal density differed among the three defined regions for cotyledons with EPF1-sectors decayed with a distance (p = 0.00028). Our findings are consistent with the role of EPF1 as secreted peptide inhibiting stomatal development and indicate that both EPF1 and STOMAGEN have limited effective range. Due to a rapidly changing heterogeneity of peptide’s impacts on stomatal density in a gradient manner, however, we conclude that precise quantification of their effective range requires a different metric.

### Spatial Autocorrelation SPACE Analysis Quantifies 2-D Spatial Patterning of Stomata

Our goal here is to quantitatively determine the effective range of peptide signals influencing epidermal patterning. However, currently available and widely-adopted quantification methods, stomatal density and index do not take into account of any 2-D spatial information, whereas they can only infer that the non-cell autonomous effects exist (Figs. 1–3). The stomatal density represents numbers of stomata in a given region of interest (ROI), ant the stomatal index represents a percentage of stomata in a given numbers of epidermal cells. With these simplistic parameters, it is not possible to normalize the inherent heterogeneity of mosaic sector size and geometry, which are constrained by individual size and geometry of epidermal cells constituting GFP_ER_ sectors. Likewise, the exact locations and numbers of individual sectors within a field of cotyledon epidermis will be unique to individual heat-shock events. Hence, it is imperative to develop a new technique for quantitative description of stomatal spatial patterning.

To this end, we adapted a statistical technique used by astrophysicists to measure spatial correlation between galaxies at different separations (Landy and Szalay, 1993; Peebles, 1974) (Fig. 4). Stomata are treated as spatial coordinates generated from an unknown probability distribution that determines their spatial patterning (Fig. 4A, D, G). Unlike the probability itself (Fig. 4C, F, I), which cannot be determined from the sample alone, the spatial correlation function can be calculated directly from the stomata as an accurate and effective approximation of the true probability distribution. The spatial correlation statistic describes this probabilistic distribution of stomata as a function of distance from a sector edge (Fig. 4). If, at a certain distance away from the edge of a GFP_ER_ sector, stomata are more likely to be found than randomly distributed, stomatal production is positively correlated with sector location (Fig. 4E, F). If stomata are less likely to be found than a randomly generated point, stomatal production is negatively correlated with sector correlation (Fig. 4G, H). If stomata production at a distance is equally likely as random point generation, this implies zero correlation between stomata production and the sector at that range (Fig. 4B, C).

**Figure 4.**
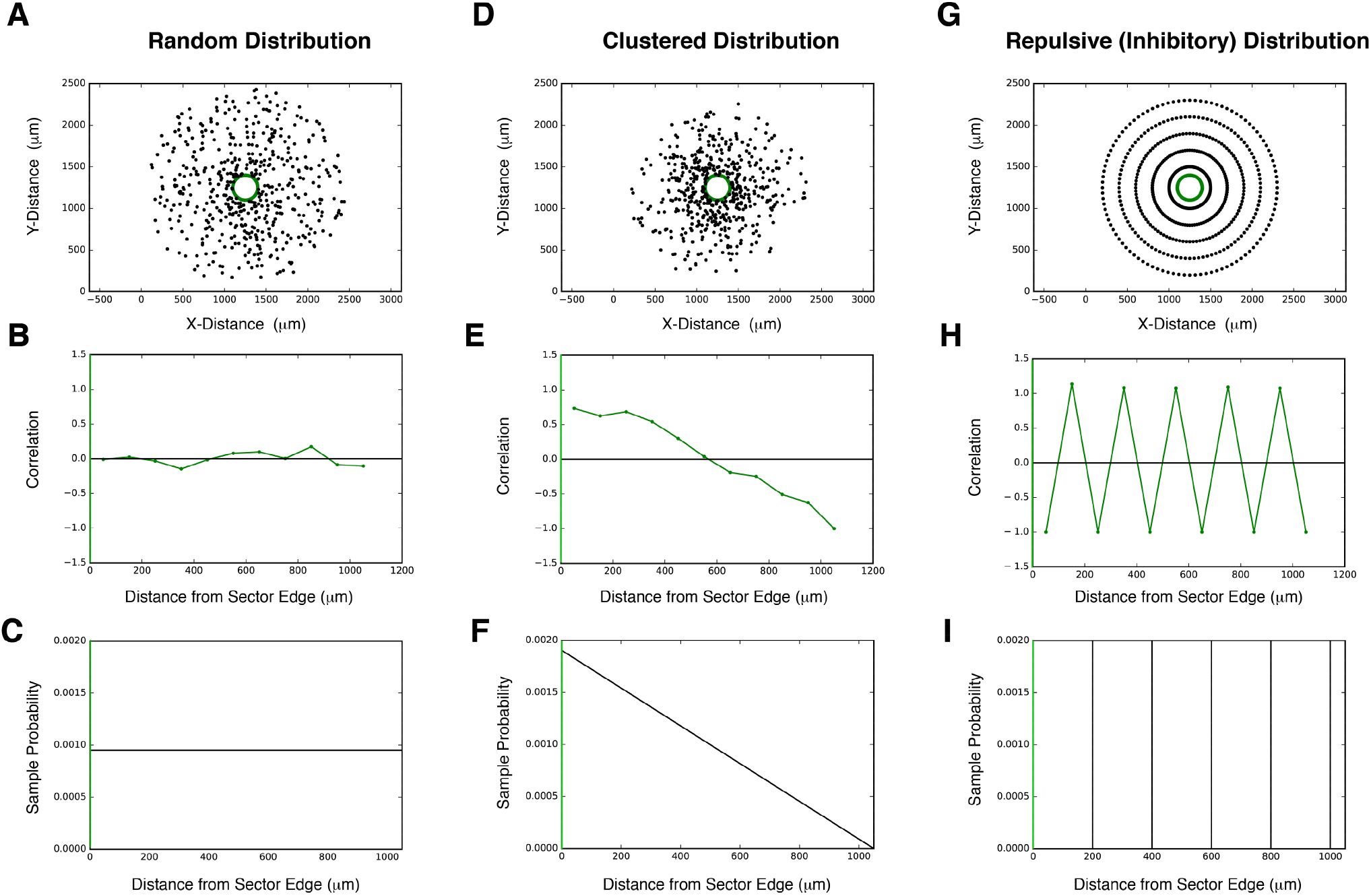
Simulating 2-D Spatial Patterning with SPACE. (A-C) Representative example of a uniformly random distribution and its statistical properties relative to a sector. Sample stomata were generated (A; black; n=500) relative to a sector (A; sector boundary highlighted in green). Stomata were generated according to the probability distribution in (C) and its stomata-sector correlation function in (B) was calculated directly from the generated points in (A) (see methods for calculation). The correlation function is close to zero both near and far from the sector, consistent with a uniformly random probability distribution. The X-axis 0 in (B, C) corresponds to a sector boundary (green). (D-F) Representative example of a clustered distribution and its statistical properties relative to a sector. Sample stomata were generated (D; black; n=500) relative to a sector (D; sector boundary highlighted in green). Stomata were generated according to the probability distribution in (F) and its stomata-sector correlation function in (E) was calculated directly from the generated points in (D) (see methods for calculation). The correlation function is highly positive at close distances, consistent with strong clustering of stomata near the sector. The X-axis 0 in (D, E) corresponds to a sector boundary (green). (G-I) Representative example of a uniformly-spaced distribution and its statistical properties relative to a sector. Sample stomata were generated (A; black; N=500) relative to a sector (A; sector boundary highlighted in green). Stomata were generated in rings around the sector, each a radius of 200 microns larger than the previous, corresponding to the probability distribution in (H). The stomata-sector correlation function in (I) was calculated directly from the generated points in (G) (see methods for calculation). As distance increases outward from the sector, the correlation function oscillates between positive and negative, corresponding to regions of stomata (positive) and empty space (negative). The X-axis 0 in (H, I) corresponds to a sector boundary (green).

To calculate spatial correlation, we plotted 2-D positions (XY coordinates) of every single stoma on an entire cotyledon, sector outlines, and cotyledon outlines from each full tile-scanned confocal Z-stack images of entire cotyledons (Fig. 5A, top and middle). To compare, we computationally generated a thousand sets of random point distributions of ‘dummy’ stomata, which are exactly the same total numbers as that of the ‘real’ stomata within the identical cotyledon outline (Fig. 5A bottom; see Methods). The nearest Euclidan distance was calculated between each stoma and the edge of a sector outline, excluding stomata inside the sector, and for every random set, the nearest Euclidean distance was calculated between each random point and the edge of a sector outline. (Fig. 5B; see Methods). After repeating this process for a thousand equally-sized sets of random points per cotyledon, the aggregate distribution of distances between random points and sectors approached its expected probability and allowed us to calculate stomatal spatial correlation as a function of distance (see Methods for further details and calculations). Because the random point sets are equal in size to the stomata index, and because the random points are generated within the same cotyledon outline as the stomata, this method enables us to quantify changes in stomatal distribution at different distances relative to GFP_ER_ sectors, independent of leaf shape, leaf size, sector placement, or sector size in a way stomatal density could not. Furthermore, the magnitude of spatial correlation quantifies the degree of change in stomatal distribution, allowing us to measure how the influence of peptide overexpression changes with distance.

**Figure 5.**
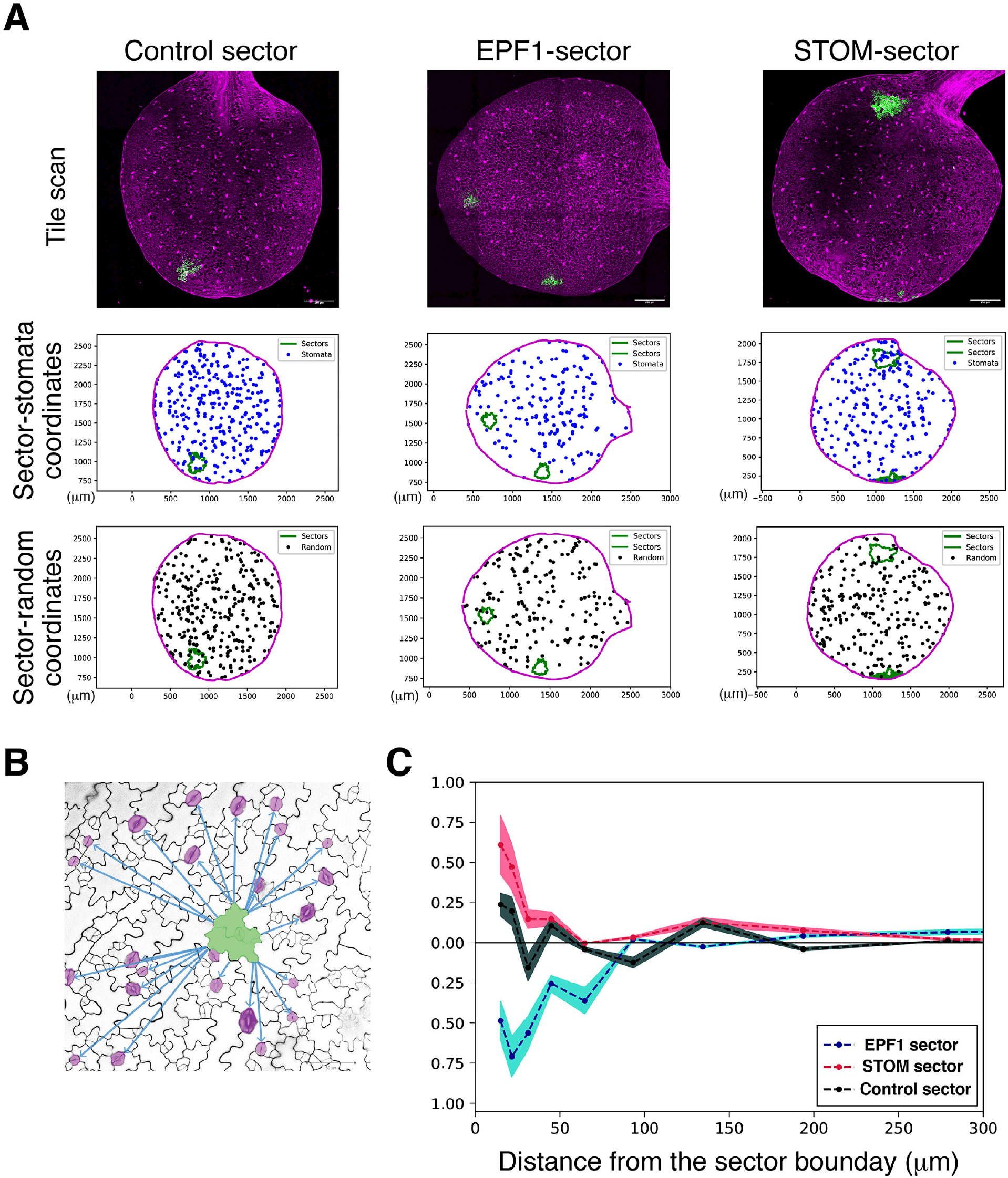
SPACE Analysis Determines the Effective Range of Signaling Peptides. (A) Representative data for SPACE pipeline. (Top) Representative fully-tiled Z-stack confocal microscopy of entire cotyledons with sectors expressing vector only control (left), EPF1 (middle), and STOMAGEN (right). Scale bars, 250 μm. (Middle) Plot of the tiled confocal images. XY-coordinates of cotyledon outlines (magenta), sector outlines (green), and all stomata on the entire cotyledon (blue) are registered. Number of stomata in each image: n= 311 (Control), 187 (EPF1), 237 (Stomagen). (Bottom) One representative plot of the 100 plots of randomly-distributed virtual stomata (black dots) with the identical n to the actually observed stomata in the images above. (B) Schematic diagram of SPACE analysis. Here, quantitative measurements were performed for the nearest distance between the edge of a sector (green) and every single stoma (magenta) as well as the nearest distance between the edge of a sector and every single random dot (randomly-placed virtual stoma) generated computationally (see panel A, middle). See methods for calculation. (C) SPACE analysis plot. The autocorrelation of sector to stomata in the function of distance from the sector boundary. Control sector autocorrelation (gray) exhibits subtle peaks at proximity ~50 μm and at ~150 μm, a latter of which may correspond to two stomata separated by one-cell spacing rule. The STOMAGEN-expressing sector (red) exhibits a strongly positive correlation at the sector boundary, which decays within the first ~60 μm. By contrast, EPF1-expressing sector (blue) exhibits a negative correlation that gradually decays at around ~160 μm. Colored area represents 95 % confidence range.

### EPF1 and Stomagen Differ in Effective Range

The SPACE analysis generated a probabilistic distribution of stomata in the function of distance from the sector boundary (Fig. 5). Control, empty-vector-only sector did not show substantial positive- or negative correlations but exhibited fluctuations within the short distance: immediate subtle drop and re-gained subtle peak at around 50 μm, with repeated pattern of subtle peak at around 150 μm, which may correspond to the one-cell spacing rule of stomata (stomata intercepted by one pavement cell)(Fig. 5C). At close distances, stomatal production was negatively correlated with EPF1 sectors, positively correlated with Stomagen sectors, and uncorrelated with control empty vector sectors (Figure 5C). At farther distances, spatial correlation between stomata production and peptide overexpression became zero for both EPF1 and Stomagen, demonstrating that our spatial correlation method can quantify what we visually observed from the tile scans. Through this approach, we were also able to identify the distance at which correlation becomes zero, which implies an effective range for the overexpression of each peptide. Sectors of EPF1 overexpression had an effective range of 170 μm and sectors of Stomagen overexpression had an effective range of 60 μm. These results suggest production of EPF1 is capable of affecting stomatal development at farther ranges than Stomagen. Furthermore, absence of discernable effects beyond 300 μm implies that local manipulation of stomatal patterning by sector overexpression of EPF-family peptides may not influence the global stomatal patterning throughout the cotyledon.

## Discussion

Members of the EPF-family of peptides regulate stomatal development at distinct stages to enforce proper spacing across the epidermis. Our study establishes the extent of EPF1 and Stomagen’s non-cell-autonomous capability to influence stomatal patterning. To address the limitations of standard phenotypic analyses, such as stomatal index and density, we created a computational pipeline SPACE to apply a correlation-based spatial point analysis and precisely quantify the peptides’ effective ranges of action.

### Correlation-Based Approaches in Stomatal Patterning

There is a growing need for quantifying stomatal patterns at higher spatial resolution, because stomatal mutants may have similar densities but different patterning as underlied by the distinct molecular mechanisms. A common statistic to address this is, in addition to the stomatal index, a count of stomata clusters that violate the one-cell spacing rule, thus extending stomatal index to clustering index or histograms to display their distributions. Our analysis of stomatal density across mosaic-cotyledons indicates that the reduction in stomatal density due to the presence of EPF1-sectors decayed with distance, and that its impact on the stomatal phenotype acted as a gradient. The rapidly changing heterogeneity of the peptide’s impact on patterning makes not only stomatal density a limited approach, but also the most common extensions to counting statistics such as the aforementioned cluster index. Thus, while we established the existence of an EPF1 effective range and a gradient in stomatal phenotype, quantifying their values with precision necessitated a different metric, the SPACE.

Statistical methods of spatial point analysis from other fields have begun to be embraced, and it is vital to find the specific metric suitable to extract the information each individual study needs. One such technique is the use of Betti numbers from Persistent Homology (Haus et al., 2018). Applied to stomatal patterning, the 0^th^ Betti number counts the number of stomata clusters (called “components”) across a leaf that remain separate when stomata are connected by a radius of given distance. Increasing the radius gradually decreases the total number of connected components, until they eventually merge into a single set, allowing one to see how the topology of stomatal distribution changes across varying spatial resolutions. To elucidate how local overproduction of stomatal peptide impacts patterning, it is necessary to utilize an algorithm that can determine the strength of disruption as a function of distance from a particular location (the sector), rather than the pattern’s overall connectedness. It would also be difficult to interpret Betti numbers for *EPF1*-overexpressing mosaics in particular, as there are little to no stomata in or near the sectors to connect to, no matter the distance. Therefore, we require our choice of metric to measure pairwise interactions and to have a bivariate form applicable to two separate point distributions: the sector outline and the stomata.

EPF1’s longer correlation length compared to Stomagen’s may come as a surprise when it is known that in wild-type, Stomagen is first expressed in the subepidermal tissue, which eventually differentiates into the mesophyll, before secreting to influence stomatal patterning in the epidermis. By contrast, EPF1 expression is in the epidermis itself. In the context of an activator-inhibitor system such as in Turing patterns, though, it is necessary that the inhibitor is longer in range than the activator as observed in this study (Kondo and Miura, 2010; Meinhardt, 2012). Regardless, before interpretation it must be noted that a peptide’s correlation or anticorrelation length, also describable as an effective range of action, is not equivalent to an effective range of diffusion. Mechanisms that contribute to an effective range of action also include the threshold of concentration each peptide must have within a cell to change a cell fate decision, the geometry of cell expansion (e.g. pavement cell geometry), and potential regulatory feedback loops. For instance, clusters of stomata produced by a Stomagen sector might produce and secrete EPF1 to cells further away, buffering against the Stomagen overexpression. Further studies are required to elucidate the degree to which these individual mechanisms contribute to the measured correlation lengths and amplitudes.

However, this approach of quantification enables each of these mechanisms to be viewed as a variable that fine-tunes the correlation function. Different features of the matter correlation function enable physicists to study the mechanism that dominates that region of the function, such as gravity or baryonic acoustic oscillations (Cole et al., 2005; Eisenstein et al., 2005). Analogously, different features of a stomatal correlation function may correspond to specific genes or mechanisms in stomatal patterning. Our SPACE pipeline is not limited to the context of stomatal development, either: It could be utilized for quantitative analyses of phenotypic characteristics and mathematical constraint broadly to the study of spatial patterns of individual cell fate, such as floral spot patterning (Ding et al., 2020).

### Local and Global Patterning in Stomatal Development

It has been reported that Stomagen as well as EPFL4 and EPFL6/CHALLAH are expressed in the non-epidermal tissues, but they could modulate stomatal patterning (Abrash et al., 2011; Kondo et al., 2010; Sugano et al., 2010; Uchida et al., 2012). Consistent with these findings, our study identified EPF1-expressing sectors exclusive to the mesophyll still inhibited stomatal development in the nearby epidermis (Fig. 2). Combined, these results highlight the necessity of viewing stomata development as a multidimensional system that acts and coordinates across multiple tissues. EPF peptides may play a key role in the inter-tissue communication between the stomata mediating gas exchange and the photosynthetic mesophyll. A recent study highlights the importance of mature functional stomata and actual gas exchange for mesophyll air-space morphogenesis (Lundgren et al., 2019). Thus, inter-tissue-layer communication involves peptide signaling at an early developmental stage and mechanical/physiological feedback during maturation. The expression of EPF peptides in internal tissues also raises the question of stomatal signaling between the abaxial and adaxial sides of the leaf. In future studies, the correlation in stomata positioning between the abaxial and adaxial stomata on the same cotyledon could be measured.

Previous studies have shown the presence of long-range hormone signaling that acts on stomatal development (Casson and Gray, 2008; Qi and Torii, 2018). In this study, we developed a pipeline that enables the quantitative measurement of spatial correlation and density at different scales of distances, separating local and global features of stomatal patterning and production. Our SPACE analysis could be used to address whether local manipulation of stomatal development may in turn influence the global stomatal patterns. For instance, locally upregulated EPF1 or Stomagen signaling could impinge on longer-range hormone signaling, such as auxin, to induce compensatory increase or decrease of stomatal development in globally. In fact, auxin and EPFL2 peptide signaling pathways constitute negative feedback during leaf morphogenesis (Tameshige et al., 2016). On the contrary, we observed that in the epidermal tissue defined as far away from Stomagen-expressing or EPF1-expressing sectors, stomatal patterning returned to normal both in density and in correlation (Figure 4). The lack of evident compensation could be explained by several possibilities. For instance, local manipulations of small EPF1/Stomagen-expressing sectors are not sufficient to trigger above-threshold compensatory response. It has been reported that overall mechanical properties of leaf epidermis could impact the polarity of a stomatal-lineage cells (Bringmann and Bergmann, 2017). Secondary changes in stomatal signaling due to sector overexpression may ‘buffer’ the global influence. Our system may be more applicable to studying the global ripple of local perturbations in mature leaves, as physiological feedback increases in importance. With the pipeline to detect and quantify local vs. global patterns in hand, future studies of mechanical and physiological feedbacks will provide the full picture of stomatal development in the context of whole functional leaf.

## MATERIALS AND METHODS

### Plant Materials and Growth Conditions

*Arabidopsis thaliana* Columbia (Col) accession was used as wild type. The Cre-Lox Gal4-UAS system used was reported previously [2]. Transgenes were generated by genetic crosses or Agrobacterium-mediated transformation (see Method Details) in the Col-0 background, with genotypes confirmed through PCR. See Table S1 for a list of plasmids generated in this study and Table S2 for a list of primer sequences used for cloning and genotyping. Seeds were sown on 0.5 × Murashige and Skoog (MS) media containing 1 × Gamborg Vitamin (Sigma), 0.75% Bacto Agar, and 1% sucrose. After stratification at 4°C for 2 days, seeds were grown in long-day condition at 21°C. To generate mosaics, seedlings 24 hours after gemination received heat-shock in a 37°C incubator as described below.

### Molecular Cloning and Generation of Transgenic Plants

The two component Cre-Lox system described previously (Heidstra et al., 2004) was modified to express full-length EPF1, EPF2, and Stomagen by the following means. EPF1 and EPF2 cDNA from pTK106 and pTK107, respectively (Lee et al., 2012), was digested with BamHl and EcoRl and ligated into pBnUASPTn to generate pTK109 and pTK110. Stomagen cDNA was PCR amplified using a plasmid pTK129 as a template and cloned into pCR2.1 TOPO vector (ThermoFisher/lnvitrogen) to generate pJS104 and sequence confirmed. Subsequently, the insert was ligated into pBnUASPTn to generate pJS105. These constructs were digested by Notl and ligated into pGll277-HSCREN2 vector to generate pTK111, pTK112, and pJS106. These three plasmids and pCB1 were individually transformed into *Agrobacterium* GV3101 (pMP90) in the presence of pSOUP (Hellens et al., 2000), and subsequently into Arabidopsis by floral dipping. More than 48 T1 plants were characterized. Three lines each of pTK111, pTK112, and pJS106 with a monogenic inheritance of selection markers were subjected to genetic crosses with the pCB1 lines, and two-to-three lines were chosen for a further analysis based on the heat-shock inducibility of *Cre* transgene as well as formation of chimeric sectors (see below). As a control, pGll277-HSCREN2 vector was transformed into pCB1 transgenic line. See Table S1 for a list of plasmids generated in this study and Table S2 for a list of primer sequences used for molecular cloning and genotyping of the transgenes.

### Heat-Shock Induction and Sector Identification

To generate mosaics, seeds after 4°C stratification were grown in long-day conditions at 21°C. 24 hours after germination, seedlings were heat-shocked in a 37°C incubator. We first tested variable duration of heat shock treatment and optimized the resulting GFP+ sector size and number. Heat-shock treatment lasted fifteen minutes to generate mosaics used in imaging experiments and lasted one hour to generate mosaics used in qRT-PCR experiments. Prior to imaging experiments for 7-day old seedlings or qRT-PCR experiments for 5-day old seedlings detailed below, seedlings with mosaic overexpression were identified using a dissecting microscope equipped with GFP fluorescence detection, Leica M165FC (Leica). We initially sought to include EPF2, an EPF/EPFL family peptide restricting the initiation of stomatal development, to our pipeline. However, due to the early events of EPF2-mediated repression of stomatal initiation during seedling germination (1-2 days)(Hara et al., 2009), a timeframe of heat-shock induced recombination and mosaic overexpression was too late to induce clear effects. For this reason, EPF2 sectors were not pursued.

### Reverse-transcription PCR

Five-day old seedlings treated with heat-shock as described above were subjected RNA preparation using RNAeasy kit (Qiagen). Subsequently, cDNA was synthesized using the iScript cDNA synthesis kit (Bio-Rad) according to instructions of the manufacturer. First-strand cDNA was diluted to a seventh in double distilled water and used as template for qRT PCR. Quantitative RT-PCR was performed as described previously (Han et al., 2018) with a CFX96 real-time PCR detection system (Bio-Rad) using iTaq SYBR Green Supermix with ROX (Bio-Rad). Relative expression was calculated by dividing *ACT2* gene expression over the specific-gene expression. For each experiment, three technical replicates were performed. RT-PCR was performed as described previously (Lee et al., 2012). See Supplementary materials Table S2 for a list of primer sequences.

### Microscopy

Confocal laser scanning microscopy images of seven-day old seedlings were taken with the Zeiss-LSM700 (Zeiss) or the Leica SP5-WLL (Leica). Cell peripheries were visualized with propidium iodide (Pl: Molecular Probes, Carlsbad, CA). GFP and Pl signals were detected with excitation at 488 nm and 555 nm, respectively, and emission at 500-524 nm and 569-652 nm, respectively. For quantitative analysis of mosaic sectors, 3-D confocal images of entire individual cotyledons (covering the adaxial epidermis and underneath mesophyll layer) were generated by tiling Z-stack frames (9-16 tiles, each with 18-45 slices for intervals of 5-10 μm covering entire cotyledon thickness and area) and stitched using Leica Application Suite AF tile scan functionality. For figure preparation, brightness and contrast of images were uniformly adjusted using Photoshop CC (Adobe).

## QUANTIFICATION AND STATISTICAL ANALYSIS

### Tile Scan Analysis and Quantification

The tile scan images were analyzed using lmaris 9.2 (Bitplane) as following. First, GFP-expressing 3-D sectors were segmented using the Surface function by thresholding the absolute intensity of the green channel, with background autofluorescence subtracted afterward. Next, sector outlines, the full cotyledon outline, and the stomatal positions were recorded with 3-D voxel spatial coordinates using the Spots function. These spatial coordinates were then exported to Microsoft Excel v.16.32 as .xlsx workbooks (Microsoft), then converted to .xlsx format for quantitative two-dimensional spatial analysis detailed below. To identify GFP sectors generated exclusively in the mesophyll, 3-D images of sectors were analyzed in the XZ and YZ planes using Leica Application Suite AF’s Orthogonal View, as well as in three spatial dimensions using Imaris 3-D volume rendering. Mesophyll-exclusisectors were segmented and their positional outlines were marked in the same procedure as vedescribed above.

### Geometric Sectors for Heat-Shocked Wild-Type

In addition to empty vector GFP sectors (“control sectors”), sector outlines were overlaid onto heat-shocked wild-type cotyledons as another control (“geometric sectors”). To generate geometric sectors, the coordinates recorded from the outlines of real mosaic GFP-expressing sectors were overlaid onto heat-shocked wild-type cotyledons. Sector outlines were transposed onto wild-type images without bias by randomly generating a shift to the center of the sector outline before plotting it on the image. If part of the newly transposed sector fell outside the boundary of the wild-type cotyledon outline, it was excluded from stomata quantification analysis. For calculating the stomatal index inside and nearby geometric sectors, any cell with at least half of its area contained inside the sector outline was considered part of the geometric sector.

### Quantitative Two-Dimensional Spatial Analysis

Spreadsheets of stomatal coordinates, sector outline coordinates, and cotyledon outline coordinates, recorded in three dimensions as described above, were processed and analyzed using SPACE (Stomata Patterning Auto Correlation on Epidermis), a pipeline of Python scripts we wrote (available at https://github.com/Toriilab/Crelox). Stomata, sectors, and cotyledons were plotted, visualized, and analyzed two dimensionally using their XY coordinates. To determine a cotyledon’s stomatal density within GFP-sectors, our script calculates the number of stomata inside a sector outline and the area enclosed by the outline. Cotyledons with multiple sectors of GFP-expression had stomata counts and sector areas aggregated before calculating the cotyledon’s overall stomatal density within sectors as single sample point.

To calculate stomatal density within a 100 μm range of sectors, a new outline was generated by applying a 100 μm radially outward shift to each sector outline coordinate. Stomata and epidermal area enclosed by the new outlines were then calculated to find the stomatal density in this region, excluding the region enclosed by the original sector outlines and any region extending beyond the cotyledon outline. For cotyledons with multiple sectors, calculations were done for the union of the new outlines, to avoid counting overlapping regions multiple times. This same process was then repeated for a 200 μm range.

To calculate the spatial correlation function between stomata positions and sector outlines, the nearest Euclidean distance was calculated between each stoma and the edge of a sector outline, excluding stomata inside a sector. The same process was repeated for a thousand times, each with independently generated random point distributions within the cotyledon outline, each equal in size to the total number of stomata across the leaf. Random point distributions were generated by first producing five times in excess the number of stomatal points within a rectangle, then running a function to keep only the random points lying within the cotyledon outline, and finally only keeping the first N points in a list, where N was the same as the number of actual stomata. Distances between sectors and stomata, and distances between sectors and random points for each independent distribution, were counted in histograms of logarithmically spaced bin widths. The spatial correlation function ζ between stomata positioning and sector location was calculated using the bivariate extension of the two-point correlation function in astronomy (Landy and Szalay, 1993; Peebles, 1974), also known as the differential form of the Ripley’s K function (Ripley, 1976):

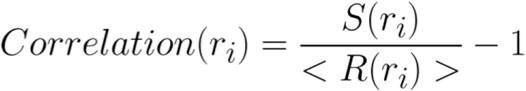

Where S(r_i_) is the number of stomata counted between a distance of r_I_ and r_i+1_ away from a sector. <R(r_i>_)> is the expected value of the number of random points counted between a distance of r_I_ and r_i+1_ away from a sector, estimated by averaging the number of points counted in that range of distance for 1000 random distributions. Because sector size may influence the range and magnitude of correlation, our code filters sectors to only analyze those within a range of area. To avoid cross-correlations on cotyledons with multiple sectors, this analysis is also filtered for sectors located within 200 μm of each other on the same cotyledon. Confidence intervals were obtained via resampling techniques.

### Stomata-Stomata Autocorrelation Function

To quantify stomatal patterning’s autocorrelation with itself, the Euclidean distance between each unique pair of stomata, for all stomata, across a cotyledon, was calculated. As described above, the same process was repeated for a thousand independently generated random point distributions within the cotyledon outline, each equal in size to the total number of stomata across the cotyledon. Distances between pairs of stomata, distances between pairs of random points within a given distribution, and distances between stomata and random points of a given distribution, were counted in histograms of logarithmically spaced bin widths. The stomata autocorrelation function was calculated using the following estimator function used in astrophysics to minimize bias and variance of the two-point galaxy autocorrelation function [6]:

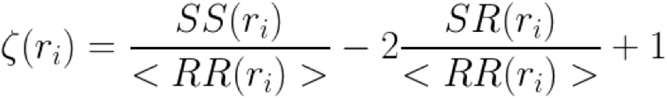

SS(r_i_) is the number of stomatal pairs counted that are separated by a distance between r_I_ and r_i+1_. RR(r_i_) is the number of random point pairs within one distribution counted that are separated by a distance between r_I_ and r_i+1_. SR(r_i_) is the number of distances separating a stomata and a random point between r_I_ and r_i+1_.All three are normalized by the total number of pairs for that variable. <> indicates expected value and is estimated by averaging across 1000 randomly generated distributions.

### Statistics

The Leica LAS AF software (Leica) and lmaris 9.2 (Bitplane) were used for image analysis as described above. Graphs were generated using R ggplot2 package or Python matplotlib. All scripts are available (GitHub account). A chi-squared test for statistical independence on 2×2 contingency tables was used for comparing the stomatal index between sectors of different peptide overexpression, in order to determine whether there was a statistically significant difference between the ratio of stomata to epidermal cells in one population versus another. The Mann-Whitney *U* test was used for comparing stomatal densities, as we do not assume stomatal density is normally distributed among the population.

## Data and Code Availability

All data, R scripts, and Python scripts for spatial analysis, are available at https://github.com/ToriiLab/CreLox.

## Acknowledgements

We thank Dr. Renze Heidstra for generously providing us the plasmids of HSpro∷CRE, 35S∷Gal4; GAL4UAS∷GFP, GAL4UAS∷Cre-Lox; Dr. Takeshi Kuroha and Janelle Sagawa for assisting molecular cloning and generating transgenic lines during the initial phase of the project; Dr. Xingyun Qi and Dr. Arvid Herrmann for expert advices on confocal microscopy; Kristen Miller and Dr. Eundeok Kim in expert assistance on qRT-PCR; and Dr. Akira Yoshinari for commenting on the manuscript. This work was supported by the Howard Hughes Medical lnstitute and Gordon and Betty Moore Foundation (GBMF-3035) to K.U.T. E.K.W.L. was supported by the Mary Gates Research Scholarship and Levinson Emerging Scholar Award from the University of Washington. K.U.T. holds the Johnson & Johnson Centennial Chair at the University of Texas at Austin and acknowledges the research support.

## Author Contributions

Conceived the project, K.U.T.; Supervised the project, K.U.T.; Designed experiments, E.K.W.L., K.U.T.; Performed research, S.Z., E.K.W.L., Analyzed data, S.Z., E.K.W.L., K.U.T.; Developed SPACE analysis, S.Z., E.K.W.L., M.F.M., B.H.; Coding, S.Z., E.K.L., B.J.H.; Writing-original draft, S.Z., K.U.T.; Writing-editing and commenting, S.Z., E.K.L., M.F.M., B.J.H., K.U.T.; Funding acquisition, K.U.T.

